# The Potential Use of Unprocessed Sample for RT-qPCR Detection of COVID-19 without an RNA Extraction Step

**DOI:** 10.1101/2020.04.06.028811

**Authors:** Arunkumar Arumugam, Season Wong

**Affiliations:** AI Biosciences, Inc., College Station, Texas 77845-5816, United States of America

**Author notes:** To whom correspondence should be addressed: Season Wong.

## Abstract

Quantitative reverse transcription polymerase chain reaction (RT-qPCR) assay is the gold standard recommended to test for acute SARS-CoV-2 infection.^1–4^ It has been used by the Centers for Disease Control and Prevention (CDC) and several other companies in their Emergency Use Authorization (EUA) assays. With many PCR-based molecular assays, an extraction step is routinely used as part of the protocol. This step can take up a significant amount of time and labor, especially if the extraction is performed manually. Long assay time, partly caused by slow sample preparation steps, has created a large backlog when testing patient samples suspected of COVID-19. Using flu and RSV clinical specimens, we have collected evidence that the RT-qPCR assay can be performed directly on patient sample material from a nasal swab immersed in virus transport medium (VTM) without an RNA extraction step. We have also used this approach to test for the direct detection of SARS-CoV-2 reference materials spiked in VTM. Our data, while preliminary, suggest that using a few microliters of these untreated samples still can lead to sensitive test results. If RNA extraction steps can be omitted without significantly affecting clinical sensitivity, the turn-around time of COVID-19 tests and the backlog we currently experience can be reduced drastically. Next, we will confirm our findings using patient samples.

## RESULTS/DISCUSSION

The sample preparation step is generally time-consuming, regardless of whether it is done manually or automated. In addition, there is a current shortage of the recommended viral RNA extraction kits needed for the Centers for Disease Control and Prevention (CDC) RT-qPCR assay to diagnose SARS-CoV-2. During a study in developing a rapid protocol for influenza (Inf) and respiratory syncytial virus (RSV) diagnostics, we investigated the feasibility of omitting the sample preparation steps to expedite the test without significantly impacting the test’s sensitivity. Using Inf and RSV clinical specimens, we successfully performed RT-qPCR reactions by simply adding a few microliters of the unprocessed sample in viral transport medium (VTM) directly into the RT-qPCR assay master mix. We then tested the approach using SARS-CoV-2 plasmid and SeraCare AccuPlex reference materials. The data presented below suggest that it is possible to skip the RNA extraction step in COVID-19 testing without a significant drop in assay sensitivity.

### RT-qPCR detection of Influenza and RSV from VTM without an RNA isolation step

We first tested the feasibility using a very small amount of sample in RT-qPCR reactions by using aerosol generating vials to spray the samples over the uncapped PCR tubes. The material in vials containing influenza A (InfA), influenza B (InfB), or RSV clinical specimens (swabs in VTM) were sprayed into PCR tubes containing primer and probe sets targeting InfA, InfB, RSV and RNaseP (RP) in master mix prior to capping the tubes and performing RT-qPCR. **Fig. 1** shows that there were positive PCR signals in the respective tubes that did not have any non-specific amplification. Though the unprocessed, sprayed samples have higher Ct values (36.5, 36.2 and 31 for InfA, InfB, and RSV, respectively) than the corresponding extracted RNA template samples (30.4, 32.6, 29.6 for InfA, InfB, and RSV). It is important to note that we were able to detect low viral load samples (Ct >30) when less than 1 μL of sample entered the tubes as measured by an analytical balance.

**Fig. 1.**
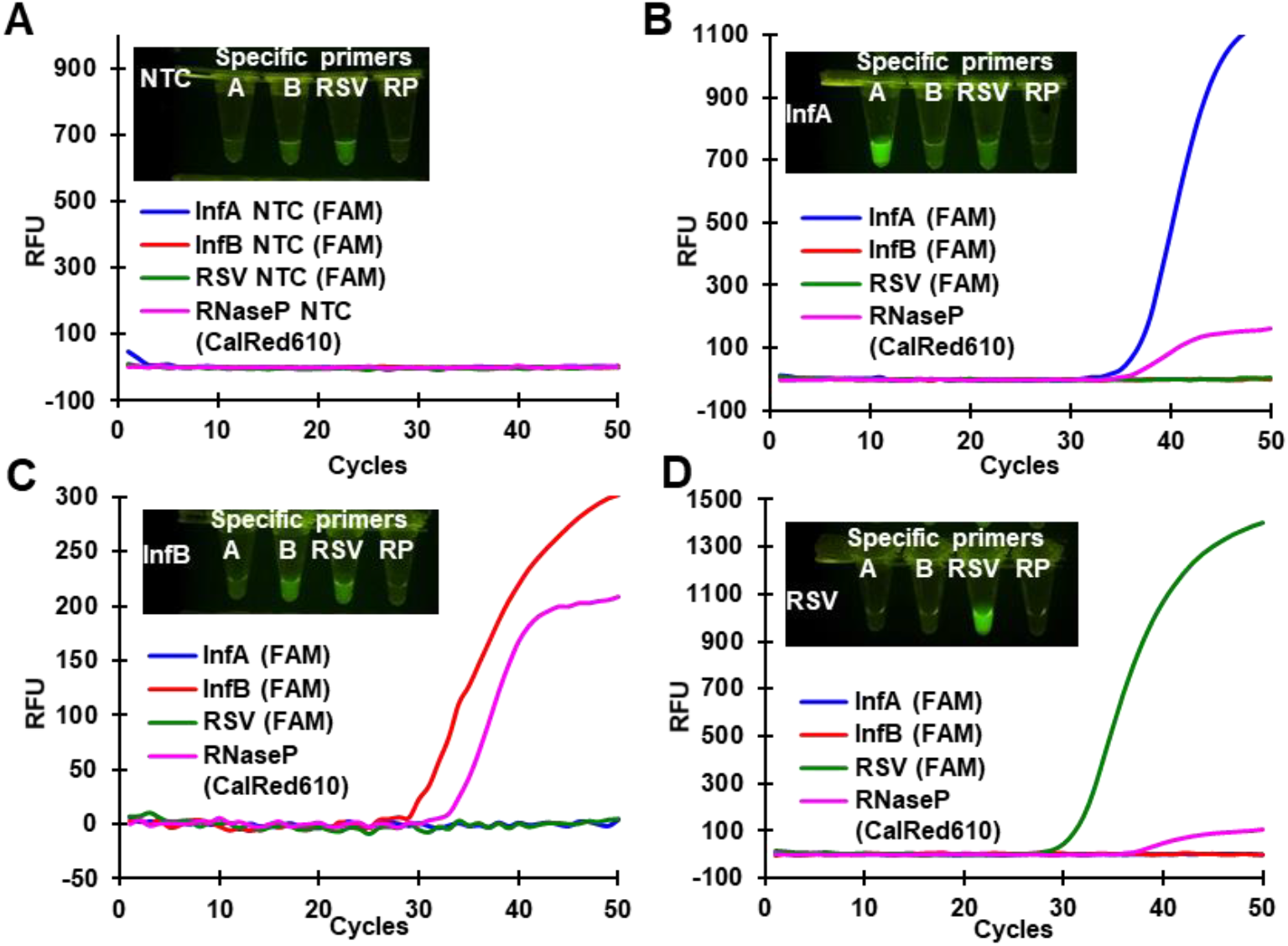
Direct detection of influenza A, B and RSV using unprocessed samples in VTM. Influenza A, B and RSV clinical specimens were diluted 1/100 times in VTM to mimic low viral load samples and sprayed into a row of four-well PCR tubes each containing InfA, InfB, RSV or RNaseP (RP) primers/probes. InfA, InfB and RSV probes were tagged with FAM dye, and the RP probe was tagged with CalfluorRed 610 (CalRed610) dye. **A** - NTC **B** - InfA clinical specimen sprayed, **C** - InfB clinical specimen sprayed, **D**-RSV clinical specimen sprayed. Inserts – Images of PCR tubes taken under blue LED illumination with an orange lens filter to capture the FAM dye’s fluorescence.

To test the extent of PCR inhibition exerted by flu specimens in VTM, we used clinical samples from Discovery Life Sciences, a Biospecimen repository. Between 0.1 μL to 6 μL of the unprocessed samples were directly spiked into the master mix with InfA primers in a 20 μL reaction mix (**Fig. 2**). An extracted template (in Promega Maxwell device) from the same sample was also amplified along with unprocessed samples. Adding more unprocessed samples (up to 6 μL) improved (reduced) the Ct values. The Ct difference between the extracted template (100 μL input and 100 μL eluate) and the unprocessed sample is minimal (29.3 vs. 30.3, respectively). Adding more than 6 μL of untreated samples has resulted in high Ct (data not shown), meaning the inhibitory effect outweighed the benefits of having more copies of the target in a reaction.

**Fig. 2.**
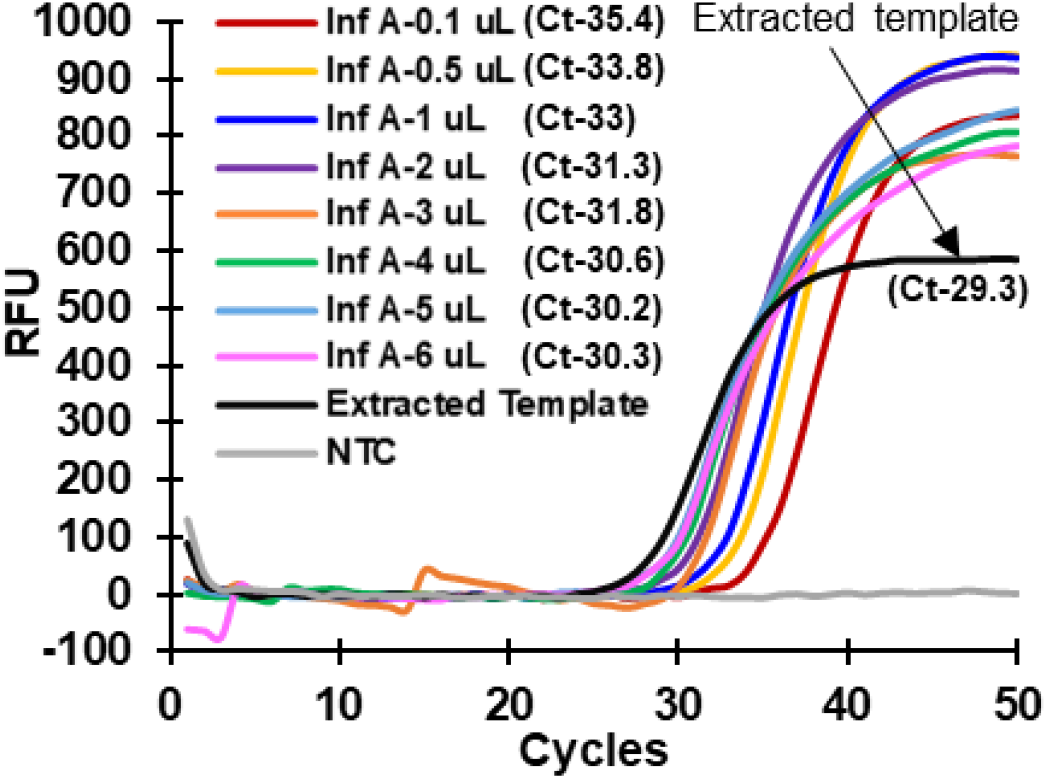
Using clinical samples without RNA extraction steps did not inhibit the reaction. In a 20-μL RT-qPCR run, 0.1 to 6.0 μL of InfA positive clinical specimens were directly added. Purified nucleic acid template (4 μL) isolated from a Promega Maxwell device was also amplified. The Ct difference between the purified template and the directly spiked clinical samples (i.e., 4 μL, 5 μL and 6 μL) is minimal.

### RT-qPCR detection of SARS-CoV-2 reference material without an RNA isolation step

We next tested whether the RNA from SARS-CoV-2 can be detected by directly spiking samples of the non-replicative recombinant virus particles (SeraCare AccuPlex SARS-CoV-2 reference material) in VTM to master mix without an extraction step. The SARS-CoV-2 virus particles were mixed with VTM to get a final concentration of 2,500 copies per mL. Different amounts (2, 4, 6 and 8 μL) of these mock clinical samples were spiked into the master mix containing CDC recommended SARS-CoV-2 RT-qPCR diagnostic panel primers N1, N2 o N3 in 20 μL PCR reactions, though we note that the N3 primers were recently removed by the CDC. Then, 100 μL of AccuPlex SARS-CoV-2 reference material was processed in Promega Maxwell device (using an AS1520 cartridge) and eluted in 100 μL of elution buffer. The extracted template (4 μL) was added to master mix, and RT-qPCR amplification was carried out in a Bio-Rad (CFX-96) thermal cycler along with the reactions with unprocessed samples.

As shown in **Fig. 3**, the SARS-CoV-2 RNA from directly spiked samples was successfully detected by the RT-qPCR reaction without a nucleic acid extraction step (N1 target shown). Up to 8 μL of VTM and AccuPlex SARS-CoV-2 mix was used in the reaction, which did not exert major inhibition on PCR amplification. The average Ct values are 38.5 (for 2 μL), 37.6 (for 4 μL), 38 (for 6 μL), and 39.1 (for 8 μL), still less than the 40 cycles that is considered to be a common cut-off for positive results. Though the theoretical copy number varied in each of the unprocessed samples (5, 10, 15 and 20 copies in 2, 4, 6 and 8 μL of input, respectively), all of them were detected. In this test, using 4 μL of the spiked sample displayed the lowest average (best) Ct. We speculate that in a 4 μL sample, the number of virus copies and PCR inhibitors in the reaction are balanced to offer maximum amplification efficiency. In comparison, the Ct value of the extracted template (20 copies/reaction) is 35.5, which is 2 cycles lower than the result from 4 μL of untreated sample (10 copies) in the reaction (**Fig. 3**). We speculate that the SeraCare matrix, containing Tris-buffered saline, glycerol, anti-microbial agents and human proteins, could be the reason for the above noticed PCR inhibition.

**Fig. 3.**
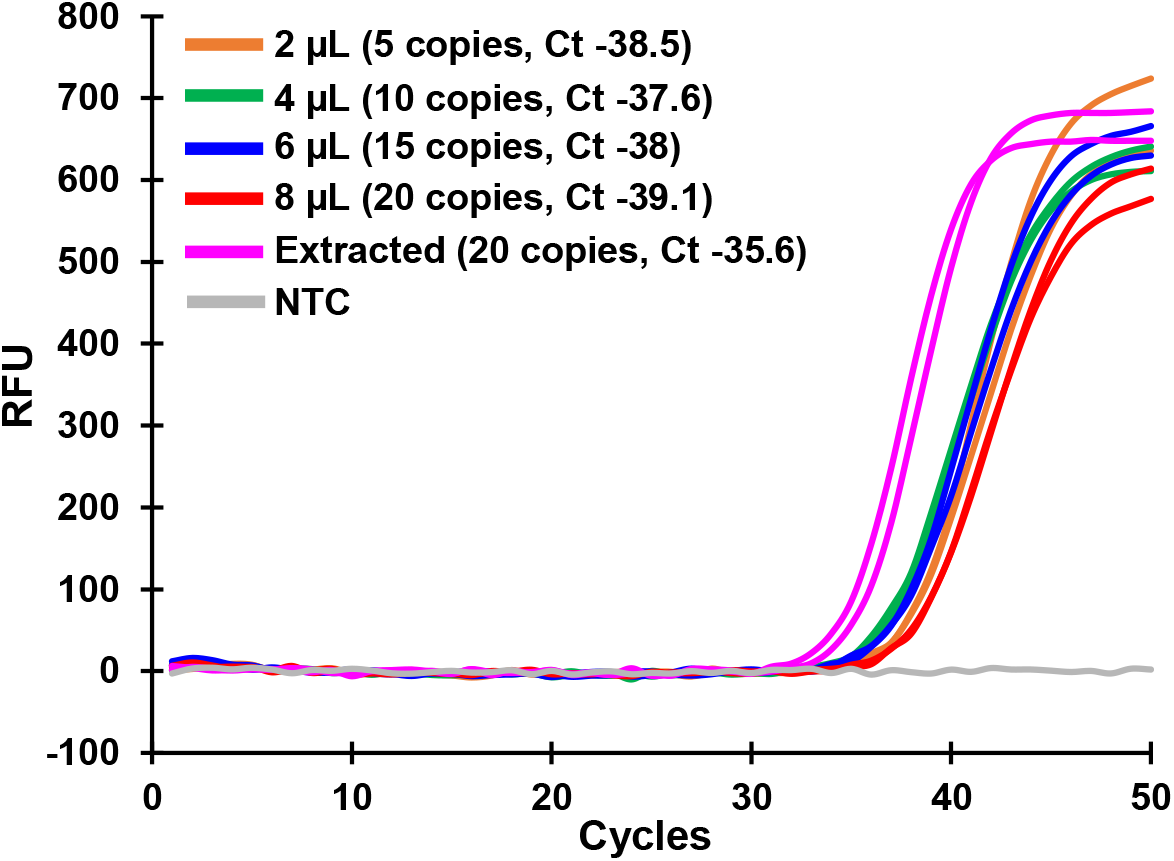
Low concentrations of SARS-CoV-2 reference material in VTM are detected without an RNA extraction step. SARS-CoV-2 non-replicative, inactivated viral particles (5,000 viral particles/mL) were diluted and mixed in an equal amount of VTM to mimic clinical samples (2500 viral particles/mL). 2 μL, 4 μL, 6 μL and 8 μL of this sample were directly spiked into a 20 μL PCR reaction mix containing SARS-CoV-2, N1 targeting primer and probe set and amplified alongside templates extracted in Promega Maxwell device (100 μL input and 100 μL eluate). RT-qPCR can detect all the directly spiked samples without any sample preparation samples (n=2).

We also tested VTM mixed SARS-CoV-2 plasmid (purchased from Integrated DNA Technologies, Inc.) using RT-qPCR. This plasmid is used as a positive control for the CDC’s SARS-CoV-2 RT-qPCR assay. The positive control plasmid was mixed with VTM and 4 μL of this mix was used for the RT-qPCR reaction. The RT-qPCR results showed that the Ct values of control (without VTM) and VTM mixed reactions (all containing 400 copies/reaction) were very similar for all three (N1, N2 and N3) SARS-CoV-2 targets (**Table 3**). We also tested 40 copies of SARS-CoV-2 plasmid using RT-qPCR, which gave a Ct value of ~38 (data not shown). This means direct spiking of a higher concentration of SARS-CoV-2 genome copies in VTM did not have a major PCR inhibitory effect in the reactions when the target concentration is high (**Table 3**). Therefore, it is likely that specimens with higher viral load could also be detected by RT-qPCR without needing an RNA extraction step.

**Table 1:**
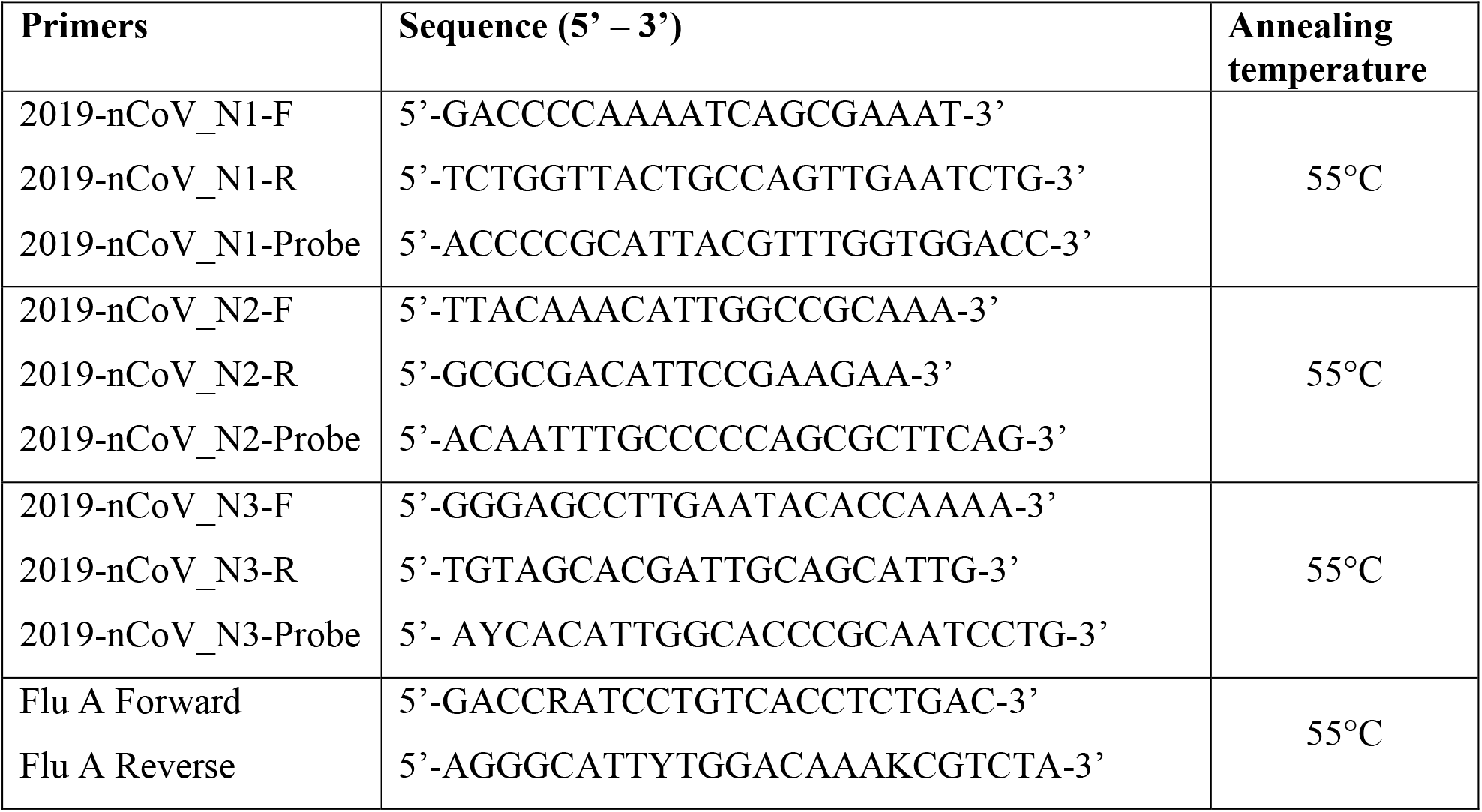
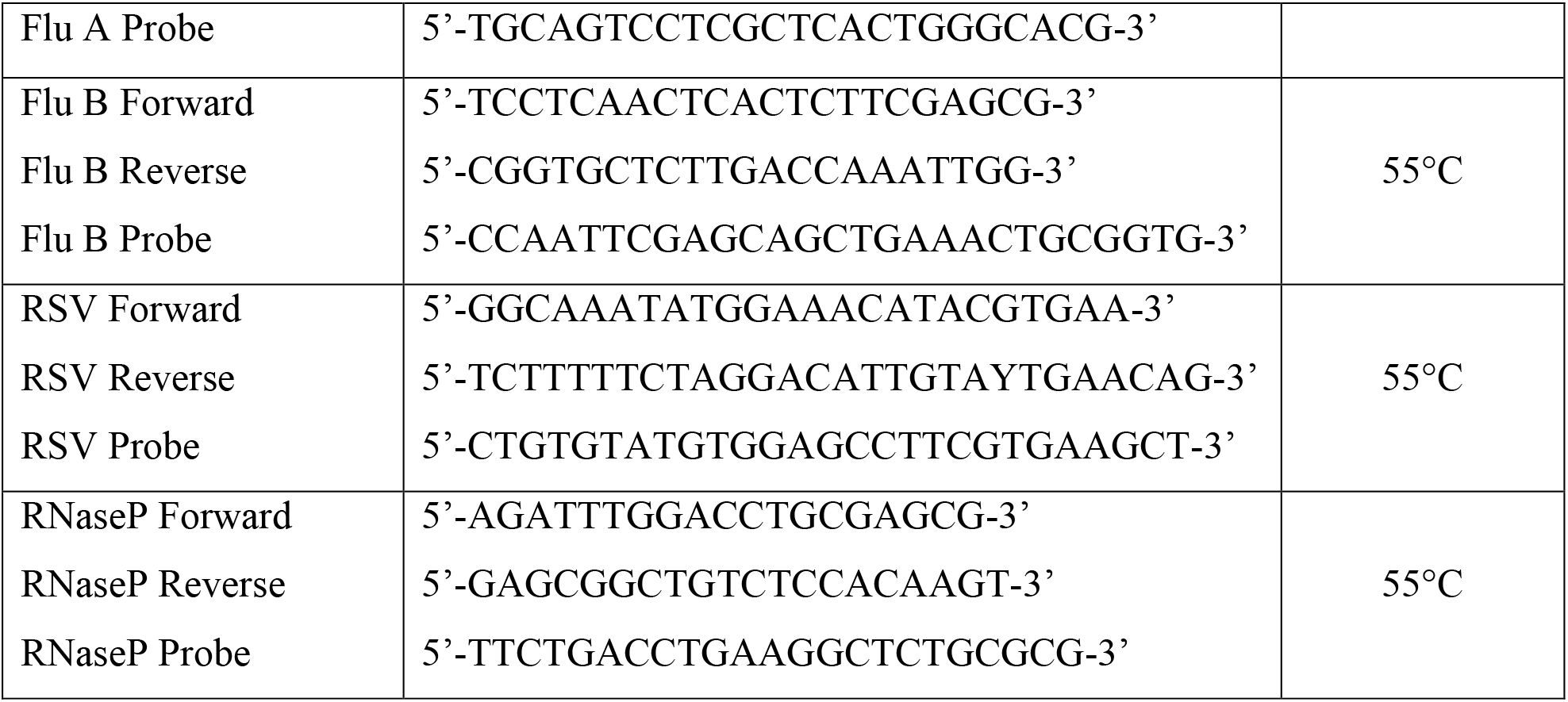
Primer and probe sequences.

**Table 2:** RT-qPCR setup.

| Process | Temperature | Time | Cycles |
| --- | --- | --- | --- |
| **UNG incubation** | 25°C | 2 minutes | 1 |
| **RT** | 50°C | 15 minutes |  |
| **Polymerase activation** | 95°C | 2 minutes |  |
| **PCR** | 95°C | 3 seconds | 50 |
|  | 55°C | 30 seconds |  |

**Table 3:** RT-qPCR detection of SARS-CoV-2 plasmid in VTM.

| Target | SARS-CoV-2 plasmid (Average Ct) | SARS-CoV-2 plasmid in VTM (Average Ct) |
| --- | --- | --- |
| N1 | 33.6 | 33.8 |
| N2 | 34.4 | 34.6 |
| N3 | 34.3 | 34.9 |

SARS-CoV-2 plasmid in VTM did not show any PCR inhibition when compared to plasmid alone. The CDC positive control plasmid for SARS-CoV-2 (from IDT) was diluted in VTM or TE buffer (control) to get 100,000 copies per mL. 4 μL of SARS-CoV-2 plasmids in VTM and in TE buffer were added to a 20 μL PCR reaction mix targeting N1, N2 and N3 genes for the detection of SARS-CoV-2. The Ct values of both the control and plasmid in VTM are very similar, indicating that the sample preparation step can be omitted in high viral load samples.

## CONCLUSIONS

Our data using both high (Ct < 35) and low (Ct >35) target concentrations suggest that since RT-qPCR is highly sensitive, using raw samples or minimal sample preparation steps might not reduce the test sensitivity as most patients tend to have a higher viral load.^5^ Also, the efficiency of extraction methods tends to drop significantly at very low target concentrations when processed through numerous washing steps. Therefore, nucleic acid loss during extraction steps may hurt the limit of detection. While further studies with patient specimens and a higher number of samples are needed to confirm our preliminary results, we report that the use of untreated samples can be a viable option during the COVID-19 pandemic.

## MATERIALS AND METHODS

### Materials and Reagents

The AccuPlex SARS-CoV-2 reference material kit (Cat. No. 0505-0126) was purchased from SeraCare (Milford, MA). Primer and probe sets for the SARS-CoV-2 RT-qPCR assay (Cat. No. 10006606) were purchased from Integrated DNA Technologies (Coralville, IA). The clinical specimens of Influenza A (DLS16-85584), Influenza B (DLS15-33890 and RSV (KH19-00715) were obtained from Discovery Life Sciences Inc. The TaqPath 1-step multiplex master mix (Cat. No. A28525) was purchased from Thermo Fisher Scientific (Waltham, MA). The viral transport media (Cat. No. R99) used for the dilution and spiking experiments was purchased from Hardy diagnostics (Santa Maria, CA).

### Preparation of low viral load clinical specimen mimics

AccuPlex SARS-CoV-2 reference material from SeraCare containing con-replicative viral particles (5,000 viral particles per mL) was mixed with equal volumes of VTM to get a final concentration of 2,500 viral particles per mL. Each microliter contained ~2.5 genomic material equivalents of SARS-CoV-2.

### Preparation of high viral load clinical specimen mimics

SARS-CoV-2 positive control plasmid (CDC recommended) was obtained from Integrated DNA Technologies (200,000 copies/μL). This was diluted to 10,000 copies/μL in TE buffer and 10 μL of this stock solution was added to 990 μL of VTM to get a final concentration of 100,000 copies/mL. A control was prepared by diluting 10 μL of plasmid stock solution in 990 μL of TE buffer. Both the VTM and TE buffer working solutions (4 μL or 400 copies in each reaction) were added into a 20 μL PCR reaction.

### Promega Maxwell extraction

The Promega Maxwell extraction was performed using AS1520 cartridge (Promega Corporation, Madison, WI). 100 μL of the sample (AccuPlex SARS-CoV-2 or Influenza A/B or RSV) was added to the cartridge and eluted in 100 μL elution buffer provided in the kit.

### RT-qPCR reaction setup

TaqPath 1-step multiplex master mix was used for the RT-qPCR reaction with specific primers and probes tagged with FAM. The final concentration of primers and probe sets for influenza A, B and RSV in the reaction is 250 nM in a 20 μL reaction.^6,7^ For the detection of SARS-CoV-2 target genes, N1, N2 and N3, the final concentration of primers are 500 nM and probes are 125 nM as per the CDC protocol^8^.

## ETHICAL STATEMENT

Specimens were purchased from Discovery Life Science. These de-identified samples are not considered to be Human Subjects.

